# Building a super-resolution fluorescence cryomicroscope

**DOI:** 10.1101/2023.11.21.567712

**Authors:** Mart G. F. Last, Lenard M. Voortman, Thomas. H. Sharp

## Abstract

Correlating super-resolution fluorescence microscopy with cryo-electron tomography is a recent advancement in the field of cryo-electron microscopy that enables targeted, high-resolution imaging of specific biomolecules of interest. Critical to this approach is that the cryo-correlated light and electron microscopy (cryoCLEM) workflow requires samples to be cryogenically fixed prior to imaging, and thus a fluorescence microscope is required that can maintain the cryogenically preserved state of the sample while also being capable of super-resolution imaging. In this report, we outline the blueprint of a microscope that was designed for single molecule localization microscopy of cryosamples, and we describe the rationale behind its design. All specifications, including a detailed 3d model of the entire assembly, are freely available via <u>ccb.lumc.nl/downloads-231</u>.

## Introduction

Correlating super-resolution fluorescence microscopy and cryo-transmission electron microscopy (cryoEM) opens up new avenues for studying biological systems at high-resolution and in native conditions^1-3^. Super-resolution fluorescence provides highly precise contrast by labeling specific components, but lacks informative content for anything other than the labelled structure. Conversely, cryoEM offers high-resolution imaging of cells in their native state, but with aspecific contrast.

To combine these two techniques, a fluorescence microscope is required that achieves a resolution high enough to be suited for correlation with cryoEM, and with which cryosamples can be imaged. In this article, we demonstrate a blueprint for such a microscope, and we outline the steps involved in constructing and operating this ‘cryoscope’. We also provide some representative data where we have used our cryoscope to perform correlative super-resolution cryo-fluorescence microscopy and cryo-electron microscopy (SR-cryoCLEM).

At a cost of only between 1-2 % that of a high-end cryoEM, and less that of commercial widefield or confocal cryo-microscope, the cryoscope can be used to locate sites of interest for cryoEM inspection with high spatial accuracy, to enrich cryoEM data by providing additional cellular context via fluorescent labelling, to screen sample thickness without requiring access to a TEM, and for numerous other applications. In our lab, the cryoscope is a cornerstone of our high-resolution cellular imaging workflow: it is routinely used for the acquisition of single molecule localization maps with ∼30 nm resolution, which are instrumental in targeting the acquisition of and identification of biomolecules within cryo-electron tomography datasets.

While there are numerous super-resolution imaging methods available that would be suited for correlation with cryoEM imaging^4-7^, the microscope described here is specifically designed for single molecule localization microscopy (SMLM)^8^, a subset of super-resolution imaging techniques comprising methods such as photo-activated localization microscopy (PALM)^9^ and stochastic optical reconstruction microscopy (STORM)^10^.

Many designs have been published for conventional ambient-temperature SMLM microscopes, and the novel requirement of maintaining samples in cryogenic conditions indeed does not change every aspect of an SMLM microscope; e.g., the image forming part of the optical system is agnostic of sample conditions. Because of the significant overlap between room-temperature SMLM microscopes and the design described here, we focus only on those features of the design and operation that are specific to cryo-microscopy, and refer the reader to previously published work for other ‘standard’ aspects^11,12^.

### Cryo-specific design considerations

#### Laser light source

SMLM methods rely on stochastic or controlled blinking of fluorescent molecules. Typically, high illumination power densities are required to achieve control over the emissive state of certain fluorescent dyes and proteins^13,14^, such as the reversibly photo-switchable green fluorescent protein rsEGFP2 (Figure 1), which is activated (converted to a fluorescent ‘on’ state) with 405 nm light and excited and de-activated (converted to a dark ‘off’ state) with 488 nm illumination. While the switching mechanisms of photo-switching fluorescent proteins are not entirely understood, particularly in cryogenic conditions^3^, it is known that photo-switching rates are significantly reduced at low temperatures^13,15^. Thus, to perform SMLM with fluorescent proteins in cryogenic conditions, multiple high power light sources are required, which are invariably laser light sources.

**Figure 1.**
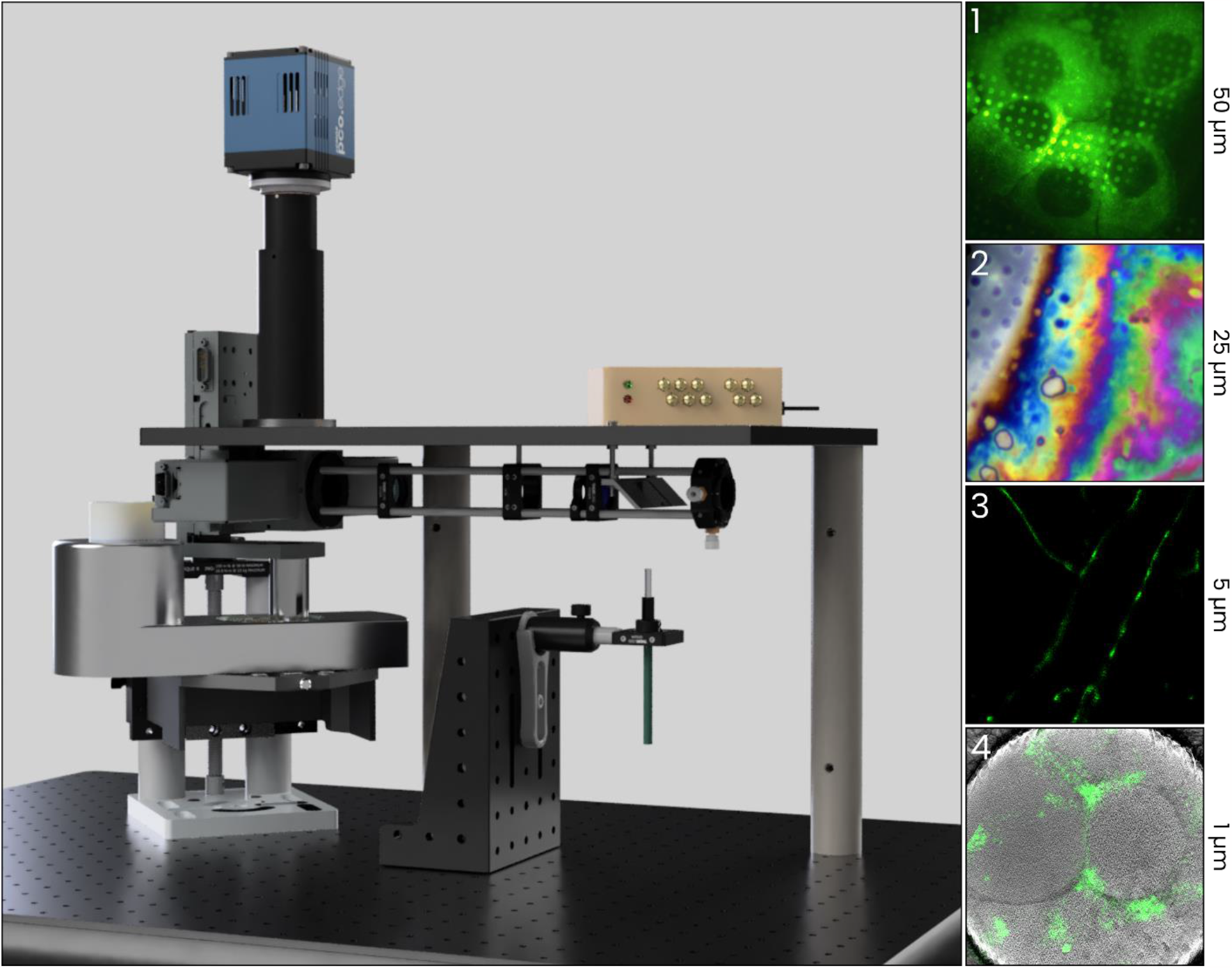
Design of the cryoscope and some exemplary results. The main image shows a 3 D model of the cryoscope – as can be seen, the optical system is relatively simple and modular. In the adjacent panels, images at various scales acquired with the cryoscope are displayed: 1) U2 OS cells expressing rsEGFP2 - IMPDH2 grown on a holey gold TEM grid (50 μm field of view), 2) an example of a three-colour composite reflected light image of a cell, which can be used to determine the sample thickness (25 μm), 3) a super-resolution image of rsEGFP2 - Vimentin (5 μm) in a U2 OS cell, and 4) a single molecule localization map of rs EGFP2-TAP1, a membrane protein of the endoplasmic reticulum, correlated with cryo - electron tomography data that reveals the colocalization of the fluorescent signal with the membranes of small vesicles.

#### Samples

High illumination power densities pose somewhat of a problem in cryogenic conditions, as the heat generated by absorption of light can lead to destruction of the sample. This issue has prompted the investigation of alternative super-resolution methods for imaging cryosamples^5,16^, and might in the future be avoided by the development of novel fluorescent probes that require less intense illumination to blink^17^. Currently, the way to avoid light-induced damage while using high illumination power densities is to select specific support grids that are capable of withstanding high illumination power densities, such as grids with low absorption support film, high mesh numbers, or grids with a low fraction of the area covered by absorbing material^18,19^.

#### LED light source

Next to the laser light source for fluorescence activation and excitation, the microscope is equipped with a multi-colour LED light source for epi-illumination. This source is not required for super-resolution fluorescence imaging, but it enables an accurate and rapid measurement of the sample thickness using a method that we have recently developed^20^ (Figure 1, panel 2). The ability to assess sample thickness in the light microscope is hugely beneficial in a correlated experiment, as it helps to determine whether a site is suited for cryoEM prior to actually taking the sample to the cryoEM.

While a multi-colour LED light source is required for thickness inspection, reflected light images acquired with any single LED source can be useful in correlating light to electron microscopy images. For most grid types, reflection contrast is mainly based on the reflectivity of the sample supporting film. In TEM images, the support film is also easily recognized. Reflection images can therefore be used as a ‘bridge’ between fluorescence and TEM images: a reflected light image contains features that are visible in the TEM, and is registered in the same coordinate system as fluorescence images.

#### Objective lens

With few exceptions, fluorescence cryo-microscopy is performed using dry lenses, i.e., without the use of immersion media. While immersion lenses can offer a higher numerical aperture (NA), technical challenges in maintaining a temperature gradient between the sample and the warmer objective lens as well as the requirement for media that remain liquid at ∼76 K render the use of immersion lenses rather impractical. It is worth noting however that cryo-microscopes using liquid and solid immersion objective lenses have been described^21,22^. Regardless, with an NA of 0.9, we routinely achieve a localization precision of 30 nm.

Besides the NA, the working distance of the objective lens is also important: sufficient spacing between the cryosample and the ambient temperature objective lens is required to ensure that the sample is not inadvertently heated by accidental contact or limited convective cooling.

Finally, the magnification of the objective lens is also an important parameter. While the magnification should be high enough to reach the Nyquist limit for the pixel size, excessive magnification can decrease the signal to noise ratio. With the high quantum efficiency and low readout and dark noise of the latest generation sCMOS detectors, in combination with the long lifetime of single-molecule events in cryogenic conditions, commonly available magnifications of 63× and 100× both meet the requirements.

In this microscope, we use only a single objective lens with a working distance of 2 mm, an NA of 0.9, and a 100× magnification. Although the addition of a second, lower magnification lens could facilitate finding areas of interest within a sample, we opt for the use of a single lens to avoid contamination of the sample that would occur during repeated switching of the lenses (see below).

#### Sample stage

The defining characteristic of a fluorescence cryo-microscope is the sample stage, which should at all times maintain the sample at low enough temperature while also providing access to an objective lens. A number of custom-built stages have been described^23,24^, but the significant investment and effort required to procure or design such stages make commercially available stages a more practical option for a group interested in using SR-cryoCLEM rather than developing it. At the time of writing, the CMS196 cryostage design by Linkam Scientific Instruments Ltd. is widely used, but there are some alternatives such as a stage by Leica that is part of a larger widefield cryo-fluorescence microscopy system.

Our design employs the Linkam CMS196v3. This is a low-profile stage that is relatively easy to integrate into a custom microscope, but also suffers from certain disadvantages – including contamination of the sample due to exposure to air, high drift, and short cold times – yet, these problems can all be mitigated by the use of flow cabinets, post-acquisition drift correction, and automated liquid nitrogen refilling devices.

We do not currently make use of a flow cabinet, with which the stage or the entire microscope could be maintained in a low humidity environment (see for example^25^).

However, a simple sleeve wrapped around the objective lens that seals the gap between the lens and the top of the stage through which humid air could flow in reduces the contamination rate. In the future, better sample stages – ideally ones that interface with cryoEM sample transport systems – might become available that are less prone to drift, require less frequent refilling, and better seal the sample compartment in order to avoid ice contamination.

### Parts list

In this section we provide an overview of the parts used in the cryoscope and discuss several alternatives. While it is possible to build a copy of our microscope with the exact parts listed below, we would recommend that prospective users investigate alternative components that are either i) better suited for their particular application; e.g., for a single-colour SMLM setup, a laser engine with a single source in combination with a spontaneously blinking fluorescent protein could be sufficient, in which case the cost can be reduced by ∼25k, or ii) improved components brought to market after publication of this book, such as alternative cryostages.

The most expensive component is the laser light engine – commercial devices typically cost tens of thousands of euros. The steep prices of certain components are well known and, although in some cases warranted (e.g. filters and detectors), in other cases large expenditures can be avoided by building custom hardware. This is particularly the case for acquisition control devices and laser light sources. As such, a number of popular open-source laser sources^26,27^ and controller^28,29^ alternatives have been developed. Whereas we make use of a commercial laser source, labs that have the required expertise and time may opt for a custom solution. For acquisition control we make use of a ‘triggerbox’; an Arduino connected to the camera, light sources, and PC, which efficiently synchronises all of the imaging hardware after it receives a user’s instructions via the microscope control software.

### Software

A number of options are available for microscope control software. Two examples are μManager, a versatile program that is widely used for diverse imaging applications^30,31^ and PYME, an open-source Python package designed specifically for SMLM^32^. During development of our cryoscope, we found that our software requirements were best met with custom software (Figure 2), which is available on our GitHub repository (github.com/bionanopatterning/cryoscope_2). Unlike e.g. PYME, this software was not written for general use, but with the limited scope of controlling our particular hardware and triggering setup. We therefore generally recommend PYME, but refer those interested in a similar custom approach to our software.

**Figure 2.**
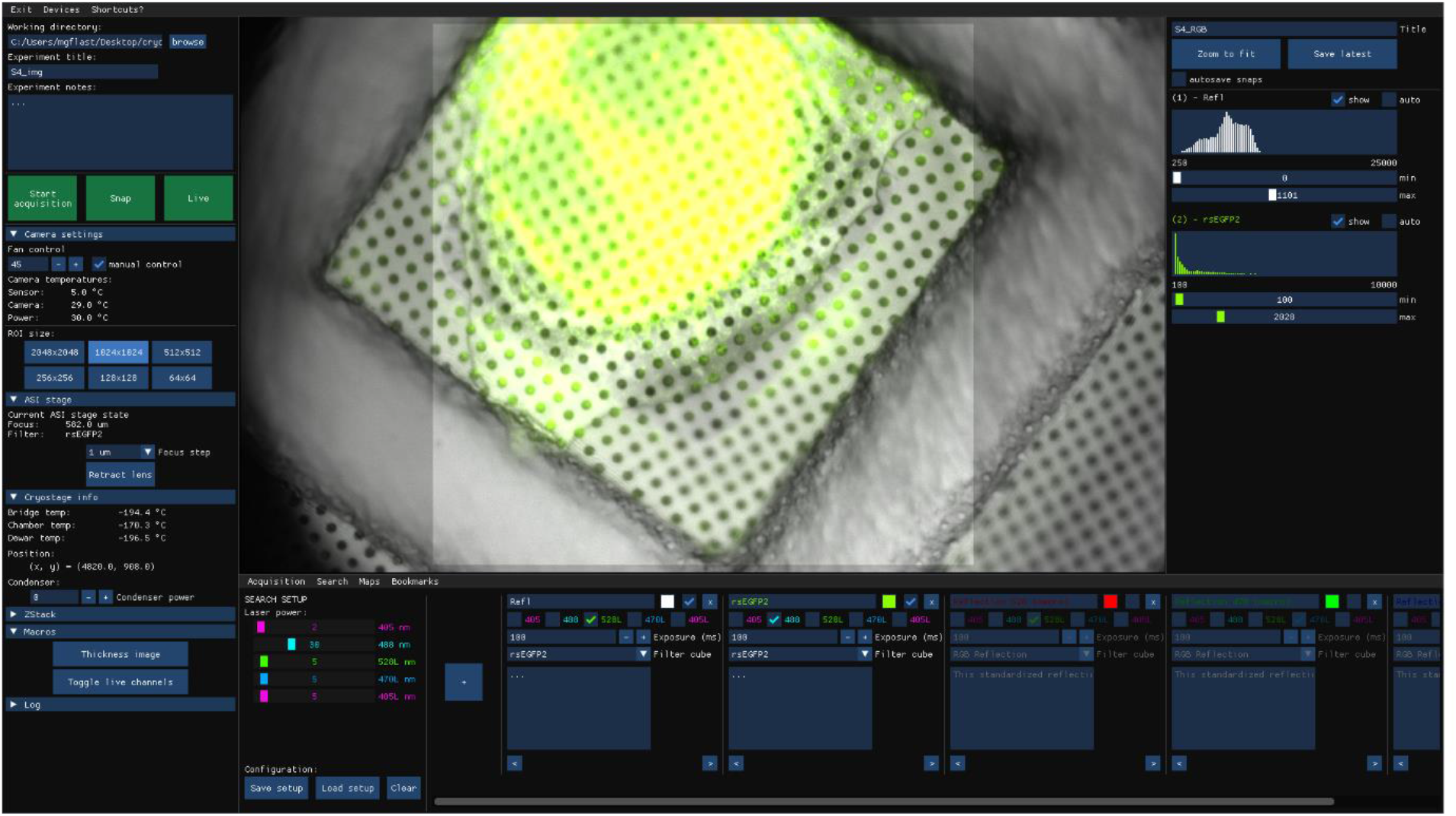
a view of the cryoscope acquisition software. In the left column, the status and configuration of the cryostage, camera, and focus and filter stage, can be inspected and controlled, and a number of other settings are available. In the horizontal bottom menu, acquisition sequences can be set up, in a minimal yet versatile interface. The column on the right allows for adjusting contrast of l ive images. The image in the centre is a composite image showing rsEGFP2 - TAP1 fluorescence (green) in an U2OS cell grown on a holey gold support film, which is highly reflective (gray)

## Results

Here we showcase an experiment that is representative of the correlated SMLM and cryoEM workflow. We used our microscope to perform SMLM on U2OS cells expressing rsEGFP2-Vimentin grown on 1.2/1.3 holey gold UltrAuFoil grids (Figure 4), but the data acquisition procedure and fluorescent label that we use (rsEGFP2) are generally applicable to any sample.

In our experience, a single-molecule localization map suited for use in identifying regions of interest for cryoEM acquisition can be acquired in only ∼5 minutes once an appropriate region of the sample is found. Finding a region of interest typically also requires a few minutes.

To create a single-molecule localization map that is suited for localizing structures of interest for subsequent cryoEM inspection, we typically use a total fluorescence imaging time of 100 seconds, e.g., 500 frames with 200 ms exposure time (Figure 4a), and an illumination power density on the order of 100–200 W/cm^2^, depending on the grid type^19^.

This is a relatively low number of frames, but it is sufficient to generate a useful SMLM image (Figure 4b,c), and since there are usually many sites of interest in one sample we currently prefer to image a larger number of sites briefly rather than extensively image a few.

Next to the fluorescence images we also expose the sample to 405 nm light in order to activate rsEGFP2 (Figure 4d) and we acquire reflection images using 528 nm LED illumination and the same filter cube as used for fluorescence imaging, which partially transmits 528 nm light (Figure 4e). These reflected light images are very useful for correlation of light to electron microscopy images.

Even though deactivation of fluorescent proteins is significantly impaired at 76 K in comparison to ambient temperature, many single-molecule events can be observed (Figure 4f-g) – particularly in thin or sparsely labelled regions of the sample: the image in Figure 4b comprises 900.000 individual localizations.

Multiple cells or regions of interest can be imaged on a single grid within a relatively short timespan; a well-trained operator can perform a SMLM imaging experiment on 5-10 sites of interest within one to two hours, including retrieving samples from storage, cooling down the sample stage, and returning samples to storage after the imaging is completed. Screening multiple grids is also facile: a grid box can be kept in the cryostage during imaging, and swapping grids from the gridbox onto the stage takes only about a minute.

Finally, SMLM imaging of cryosamples is highly compatible with cryoEM imaging. Provided illumination power densities are kept in check, samples are not devitrified by fluorescence imaging, and prolonged high-power laser illumination has also been found not to affect achievable resolution in single-particle analysis protein structure determination^33^. We believe that the combination of cryoEM and cryoSMLM is a very promising method: it allows for the localization of specifically labelled proteins of interest within the context of high-resolution electron microscopy images, thus enriching cryoEM data and enabling accurately targeted cryoEM data acquisition, which could vastly increase throughput of cryoEM imaging of rare, transient or hard to identify structures.

## Construction

The design of the cryoscope is largely modular and simple, in the sense that the number of components is low and assembly is straightforward. The triggerbox, mentioned above, provides an interface to connect the light sources, camera, and the PC (Figure 3A). We use a breadboard mounted on solid steel posts on a vibration-reducing optical table as a construction platform (label B in Figure 3). The laser illumination, LED illumination, and imaging arms of the optical path (labels C, D, E, respectively) can be assembled in isolation and are subsequently attached to the construction platform via a simple connecting interface. The focusing and filter stage (F) is a single off-the-shelf component. Finally, the cryostage is mounted in a holder that allows for careful alignment of the stage position, which in turn is carried by a manual vertical positioning stage with which the stage can be engaged to the microscope.

**Figure 3.**
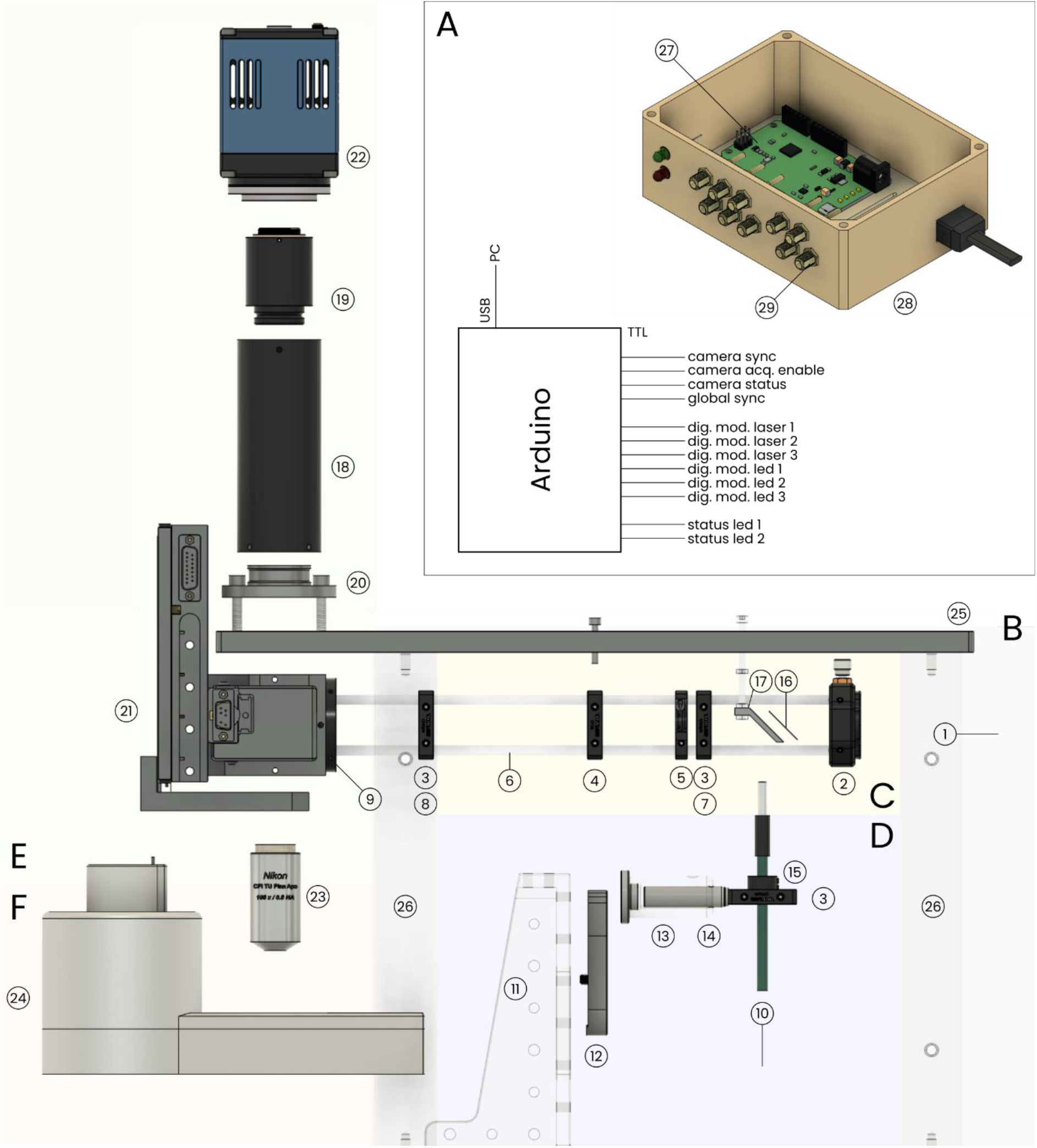
annotated exploded view of the cryoscope. Number tags indicate items listed in Table 1. Background colours as well as labels indicate modules: the triggerbox (A, white), construction platform (B, gray), excitation light path (C, yellow), LED illumination path (D, blue), imaging path (E, green), and the cryostage (F, red).

**Figure 4.**
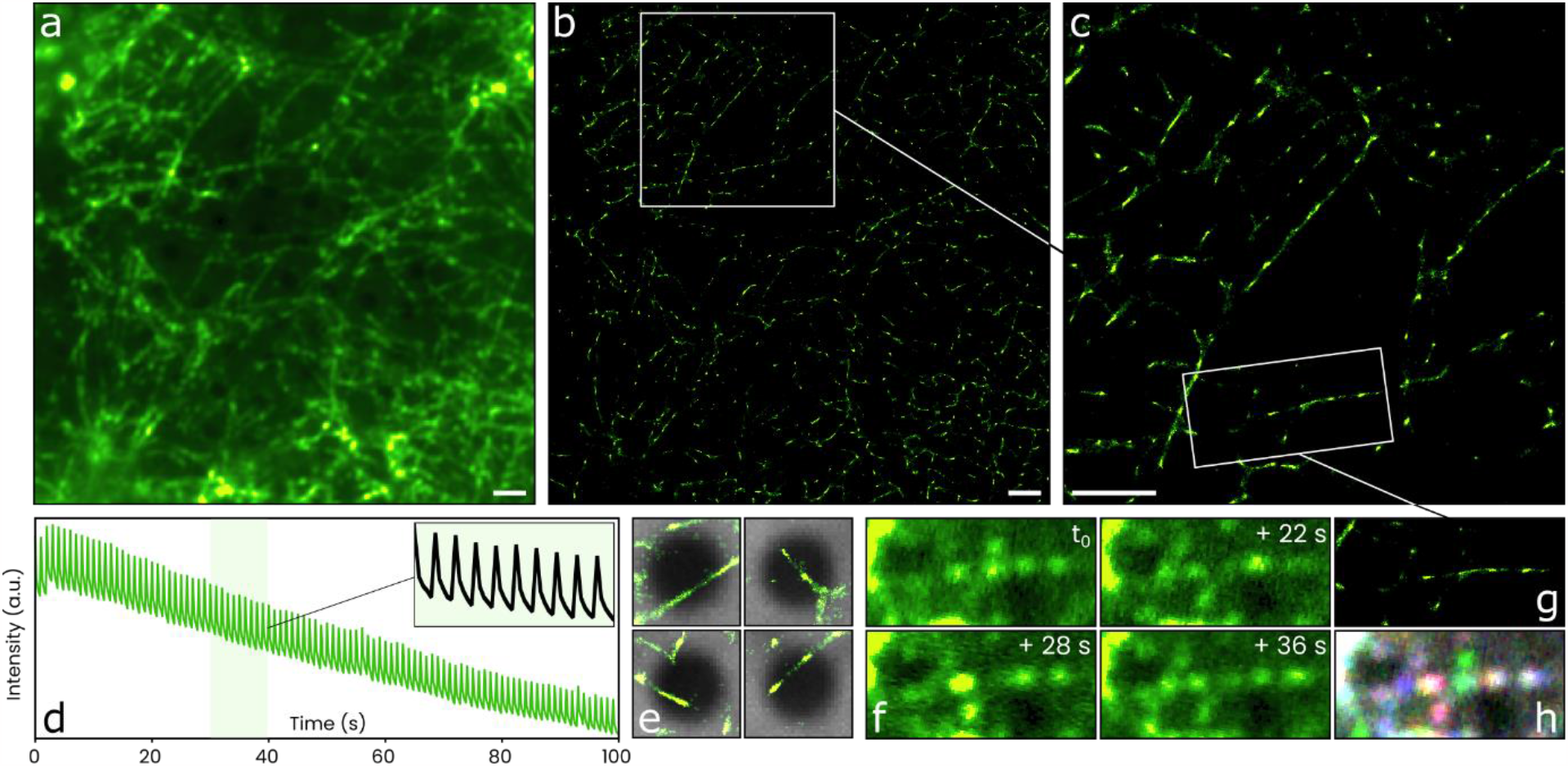
cryo-SMLM imaging of Vimentin, labelled with rs EGFP2, in a U2 OS cell. a) Sum projection image of a 500 frame timelapse video of rs EGFP2 fluorescence. To activate the fluorescent protein, a brief 405 nm illumination pulse was applied after every 5 fluorescence frames. b) Single-molecule localization map (900, 000 localizations) generated using the same dataset. c) Magnified section of the region indicated by the white square in panel b. d) A plot of mean fluorescence intensity per frame versus time since the start of the acquisition. After every 405 nm pulse, the overall fluorescence intensity of the frames increases as many molecules of rs EGFP2 are activated, and then decreases again as fluorescence is excited by 488 nm illumination, which also deactivates rs EGFP2. The inset shows a section of the graph magnifi ed. e) Examples of regions that could be interesting for cryo EM data acquisition. Vimetin fi laments can be recognized in the localization map (green), which is overlayed on reflected l ight images (gray) that show holes (1. 2 μm) in the holey gold sample sup port fi lm. f) Single frames from the timelapse dataset, showing the same region of the sample at different moments in time. Where a fi lament is seen in the final reconstruction (see panel g), fluorescence from individual molecules can be seen in the frames. g) The region of the single-molecule localization map corresponding to the frames shown in panel f. h) Three of the images in panel f combined into a composite RGB image, to visualize the relative location of the single molecule fluorescence spots. Scale bars are 2 μm.

A number of parts require custom machining: 1) the focus and filter stage is not originally intended for mounting to a breadboard, and was modified for us by the supplier, 2) a hole must be cut in the breadboard, as the imaging path runs through it – this modification can also be requested from the supplier, 3) the tube lens is not designed for mounting to a breadboard and a custom adapter is required for this purpose, and 4) a custom dovetail is required to connect the cryostage mount to the vertical translation platform. These latter two parts are simple enough to fabricate that either a supplier or a local workshop should be able to create it.

**Table 1.** list of parts used in the construction of the cryoscope. The list is subdivided into modules (blue headers), some of which (e. g. the LED illumination module) are not absolutely required for SMLM. Suppliers and supplier i tem IDs are indicated, as well as the approximate cost (€) of the part. This approximation is the most inaccurate for the more expensive parts: l ight sources, cameras, fi l ters, and objective lenses vary in price depending on the chosen supplier and specifications. Shipping and customs fees, which can be significant, are not included. The rightmost column, ‘ID’, l ists the labels used in Figure 3.

| Description | Supplier | Item ID | Cost (€) | ID |
| --- | --- | --- | --- | --- |
| **Laser illumination module** |  |  | ~50600 |  |
| Laser light engine w. SM fiber with sources:<br>405 nm / 100 mW<br>488 nm / 120 mW<br>561 nm / 150 mW | Omicron-Laserage | LightHUB | 50000 | 1 |
| XY adjustable fiber mount + fiber adapter | Thorlabs | CXY1 | 200 | 2 |
| Lens mounts | Thorlabs | CP33M | 17 x 2 | 3 |
| Assembly mount | Thorlabs | CP36M | 22 | 4 |
| Adjustable diaphragm field stop | Thorlabs | CP20D | 102 | 5 |
| Cage rods | Thorlabs | ER12 | 17 x 4 | 6 |
| Collection lens | Thorlabs | LBF254-075 | 50 | 7 |
| Field lens | Thorlabs | LA4874-A | 100 | 8 |
| Cage system attachment plate | ASI | C60-30CRM-30LM | 100 | 9 |
| **LED illumination module** |  |  | ~15600 |  |
| LED light source w. LLG with sources:<br>405 nm, 470 nm, 532 nm | Omicron-Laserage | LedHUB | 15000 | 10 |
| Various mounts | Thorlabs | AP90RLM | 191 | 11 |
|  | Thorlabs | CF175C | 14 | 12 |
|  | Thorlabs | PH50E | 28 | 13 |
|  | Thorlabs | TR50 | 6 | 14 |
|  | Thorlabs | CP33M | 17 | 3 |
|  | Thorlabs | AD11F | 33 | 15 |
| Beam combining mirror |  | 24 x 32 mm cover slip | 0 | 16 |
| Beam combiner mount | Custom made |  | 0 | 17 |
| Reflection imaging beamsplitter | Thorlabs | BSW26R | 306 |  |
| **Imaging** |  |  | ~35000 |  |
| Tube lens | ASI | C60-TUBE-B | 1000 | 18 |
| C-Mount adapter | ASI | C60-3060-C-MOUNT | 225 | 19 |
| Tube lens to breadboard adapter | Custom made |  | 150 | 20 |
| Focus and filter stage | ASI | MIM4 Slider Nikon | 6000 | 21 |
| Focus and filter stage controller | ASI | MS1+1 | 4000 |  |
| Camera | pco | edge 4.2 bi wc | 17000 | 22 |
| Objective lens | Nikon | TU Plan Apo 100/0.9 | 6500 | 23 |
| Sample stage |  |  | ~22200 |  |
| Cryostage | Linkam Scientific | CMS196v3 | 20000 | 24 |
| Vertical translation manual stage | Thorlabs | VAP4_M | 800 |  |
| Cryostage mount | Zeiss |  | 1300 |  |
| Cryostage mount adapter | Custom made |  | 100 |  |
| Filters |  |  | ~2000 |  |
| BFP channel – emission | Thorlabs | MF460–60 | 255 |  |
| BFP channel – dichroic | Thorlabs | MD416 | 277 |  |
| GFP channel – emission | Semrock | FF01-525/50 | 350 |  |
| GFP channel – dichroic | Semrock | FF495–Di03 | 390 |  |
| RFP channel – emission | Semrock | FF01-630/92 | 400 |  |
| RFP channel – dichroic | Semrock | FF570–Di01 | 280 |  |
| Mounting |  |  | ~3500 |  |
| Optical table – frame | Thorlabs | SDP6090 | 1828 | 30 |
| Optical table – breadboard | Thorlabs | B6090 | 1259 | 31 |
| Microscope breadboard | Thorlabs | MB3045 | 195 | 25 |
| Microscope breadboard support | Thorlabs | P300M | 96 x 4 | 26 |
| Acquisition control |  |  | ~2300 |  |
| PC + accessories |  |  | 2000 |  |
| Arduino Leonardo |  |  | 30 | 27 |
| Arduino Housing | Custom made |  | 0 | 28 |
| Cables and connectors | Dependent on camera and light sources |  | 10 x 25 | 29 |
| Approximate total cost |  |  | €131000 |  |

Finally, we use a 3D-printed housing for the triggerbox and to hold the beam splitter that combines the laser and LED illumination paths. Since the LED source is used for reflection imaging, which does not require bright illumination, a thin, low-reflectivity mirror is optimal for combining the laser and LED sources: it transmits most of the laser light, but also reflects a low fraction of the LED source. In our experience, a simple glass cover slip is ideally suited for this purpose.

The images below provide detailed views and additional information on each of the modules used in the construction of the cryoscope. 3D models of the full design can be downloaded at <u>ccb.lumc.nl/downloads-231</u>.

### Mounting

**Figure 4.**
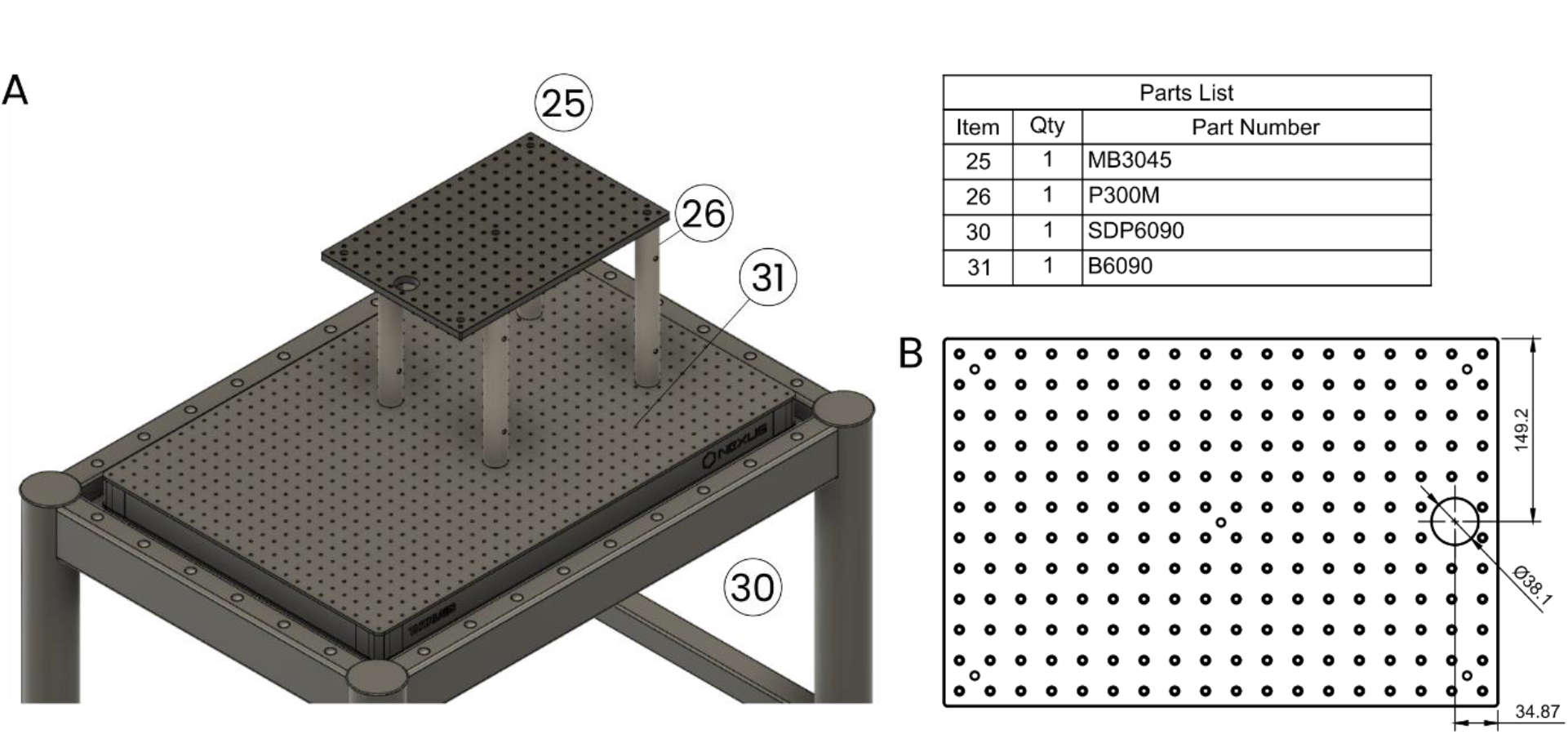
A view of the mounting platform onto which the cryoscope is assembled. A) 3D view of the mounting platform assembled on the optical table. Parts are tagged by numbers and listed in the adjacent table. B) A drawing specifying the required modifications to the breadboard.

### Laser illumination path

**Figure 5.**
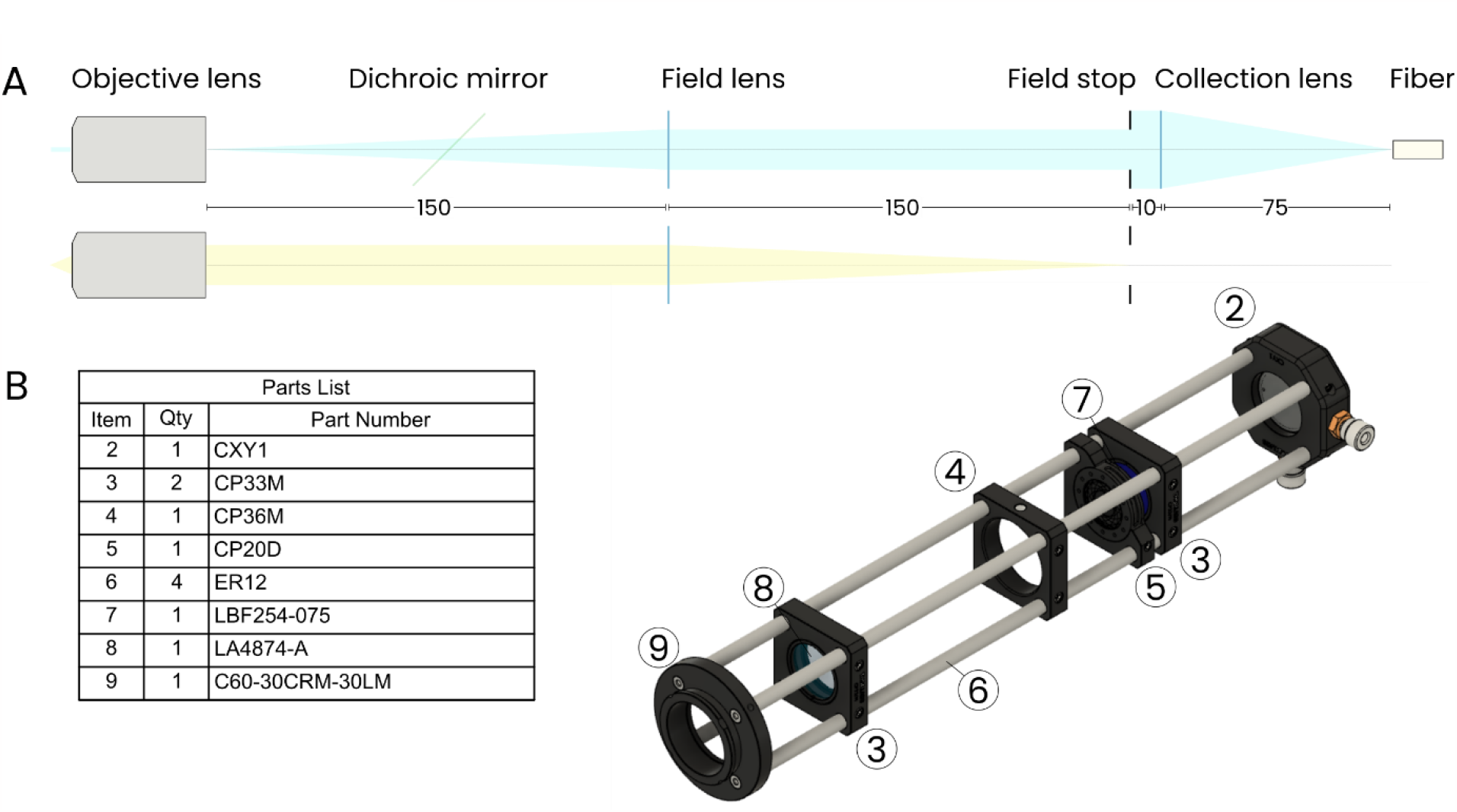
The laser illumination path. A) Two schematics representing the path traversed by laser light (top), and the positioning of the field lens and field stop (bottom). B) 3D model of the laser illumination path, with number tags corresponding to items listed in the adjacent table.

### Imaging path

**Figure 6.**
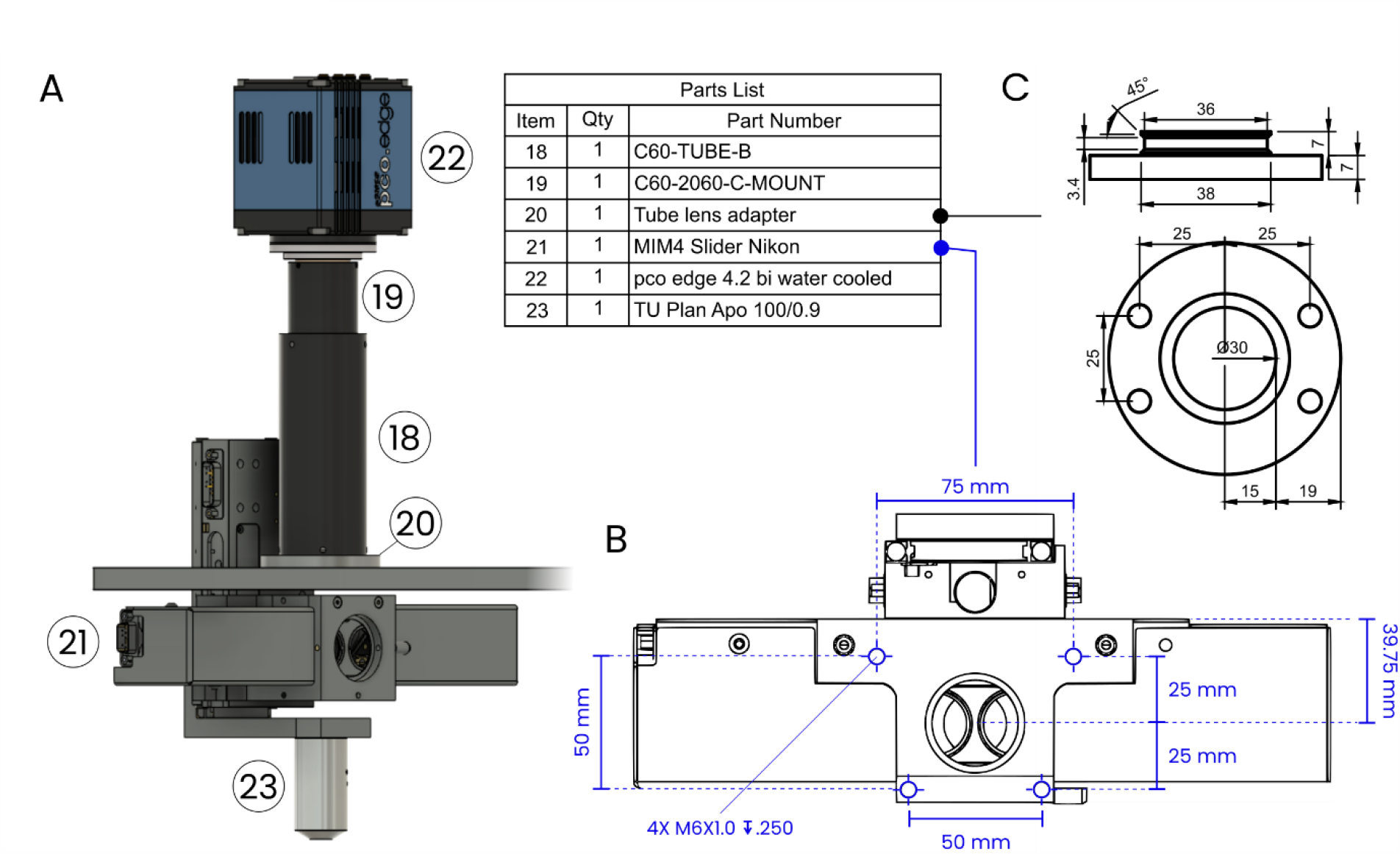
The imaging path. A) A view of the imaging path, comprising the camera, objective lens, and the combined focus and filter cube stage. Items corresponding to the indicated numbers are listed in the adjacent table. B) A drawing specifying the required modifications to the filter and focus stage. Four M6 threaded holes are drilled in the top to allow for mounting to the underside of the assembly breadboard. The supplier (Applied Scientific Instrumentation) modified this part for us; refer to the C60 - SLIDER-LUMC modification. C) A drawing specifying the design of the custom adapter that we use to mount the tube lens to the assembly breadboard.

### Cryostage

**Figure 7.**
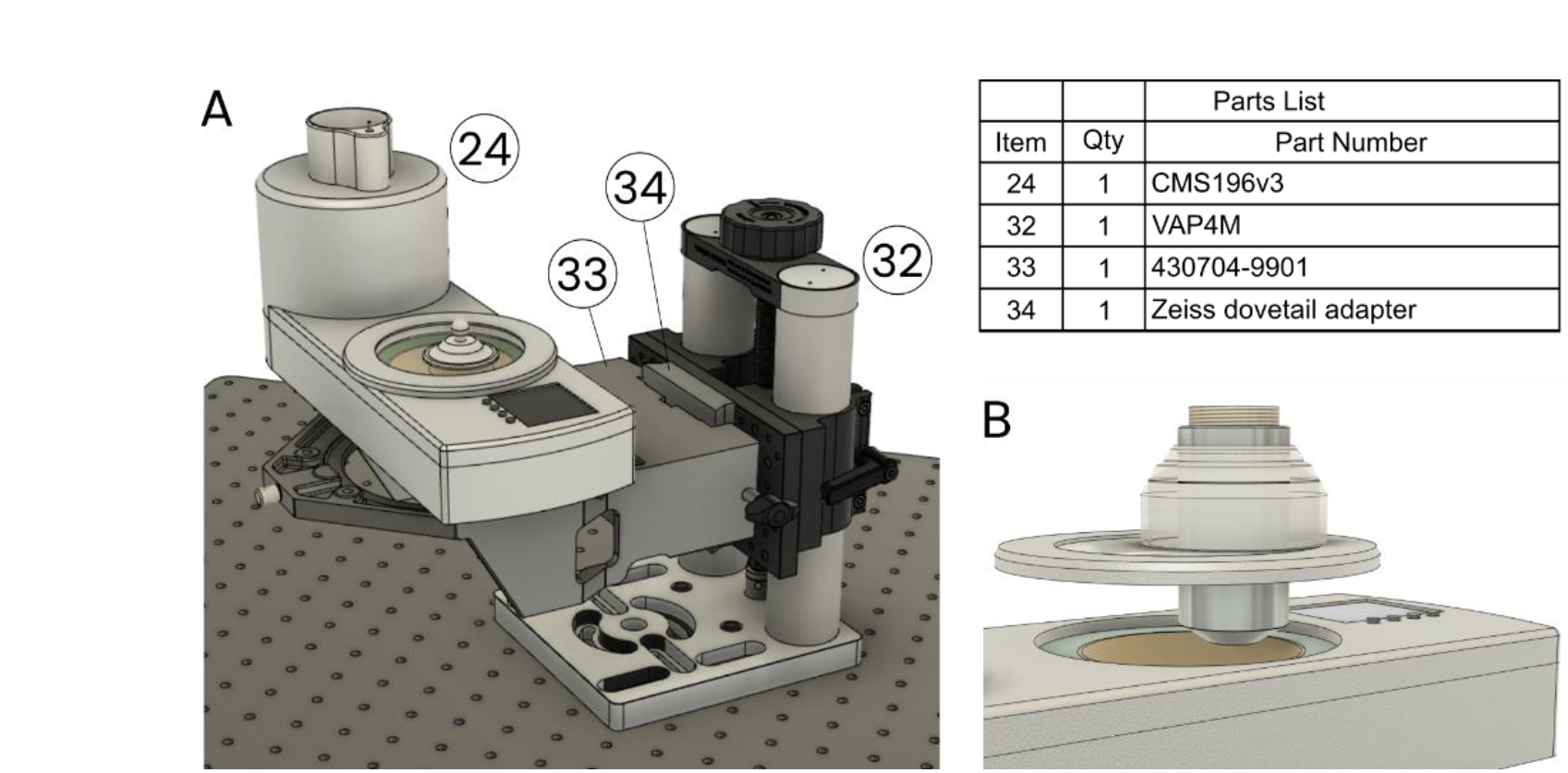
The positioning of the cryostage. A) A view of the cryostage mounted on the cryostage holder (Zeiss), which is in turn mounted onto a manual vertical positioning stage that is used to engage the stage to the microscope. B) A close - up of the cryostage with the objective lens and a protective sleeve inserted through the lid of the stage. During operation the l id covers the sample chamber – in the image here, the lid is offset vertically for illustrative purpose.

### LED illumination path

**Figure 8.**
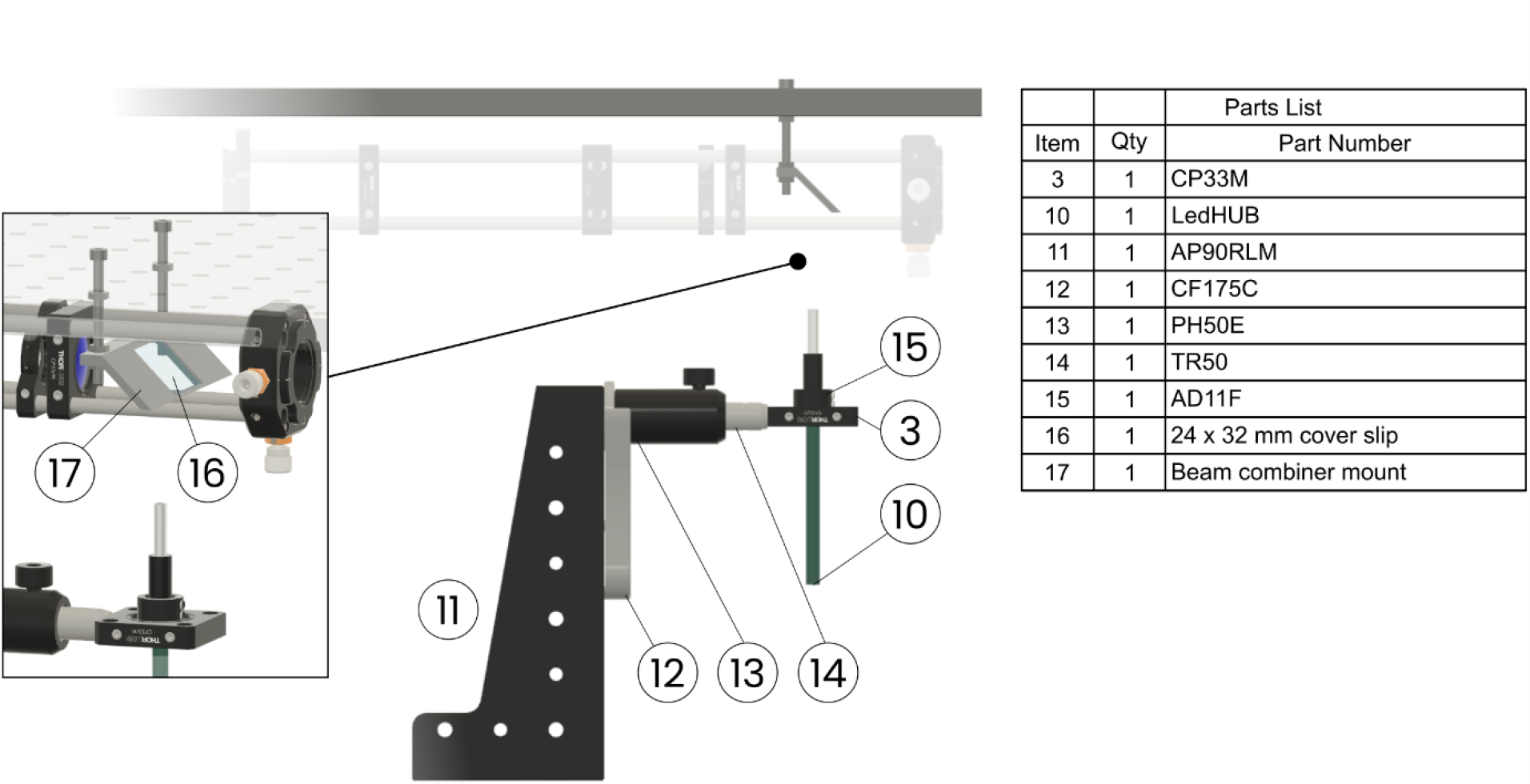
Detailed view of the LED illumination module. After constructing the cryoscope without the LED illumination module, this module can be retrofitted into the microscope. The base (parts 3, 10 - 15) is simply placed on the optical table and the beam combining mirror is mounted in a 3D printed holder that can be inserted into the laser illumination path (see inset). Part names are listed in the adjacent table.

## Conclusions

In the above, we’ve outlined the design of a microscope suited for cryo-single molecule localization imaging and demonstrated its use by imaging rsEGFP2-Vimentin expressing U2OS that were grown on grids for cryoEM imaging. One acquisition, which requires only ∼5 minutes of imaging time, can result in hundreds of thousands of single-molecule localizations across an area of ∼100 μm^2^. Within this area, one can typically identify numerous sites of interest that are also suited for cryo-electron tomography imaging. The combination of super-resolution fluorescence microscopy and cryoEM is thus a feasible and, in our experience, even a routine method that can be to accurately locate sites of interest for cryoEM and enrich cryoEM data with the context provided by single-molecule localization maps.

Future developments in hardware (e.g. new cryostages and cryo-sample transfer systems), software (e.g. for correlation), and methodology (e.g. novel fluorophores with enhanced switching characteristics) will help improve localization accuracy, sample cleanliness, imaging throughput, and many other aspects of correlated cryo-microscopy, and we are optimistic that the advantages offered by the combination of these two high-resolution imaging methods will be recognized by and accessible to increasingly more researchers. To this we hope that our design can make a contribution, and we are therefore open to communications about adaption and further development of our and similar microscope designs.

## Acknowledgements

We thank Paul Verkade for their helpful feedback.

